# Augmented gut hormone response to feeding in older adults exhibiting low appetite

**DOI:** 10.1101/2023.12.29.573652

**Authors:** Aygul Dagbasi, Jordan Warner, Victoria Catterall, Daniel R Crabtree, Bernadette Carroll, Gary Frost, Adrian Holliday

## Abstract

Age-related changes in gut hormones may play a role in anorexia of ageing. The aim of this study was to determine concentrations of ghrelin, PYY, and GLP-1 in older adults exhibiting an anorexia of ageing phenotype. Thirteen older adults with healthy appetite (OA-HA; 8f, 75±7 years, 26.0±3.2 kg·m^-2^), fifteen older adults with low appetite (OA-LA; 10f, 72±7 years, 23.6±3.1 kg·m^-2^), and twelve young adults (YA; 6f, 22±2 years, 24.4±2.0 kg·m^-2^) completed the study. Healthy appetite and low appetite were determined based on BMI, habitual energy intake, self-reported appetite, and laboratory-assessed *ad libitum* lunch intake. Participants provided a fasted measure of subjective appetite and blood sample (0 minutes) before consuming a standardised breakfast (450 kcal). Appetite was measured every 30 minutes for 240 minutes and blood was sampled at 30, 60, 90, 120, 180 and 240 minutes. At 240 minutes, an *ad libitum* lunch meal was consumed. Relative energy intake at lunch (expressed as percentage of estimated total energy requirement) was lower for OA-LA (19.8±7.7%) compared with YA (41.5±9.2%, *p*<0.001) and OA-HA (37.3±10.0%, *p*<0.001). Ghrelin suppression was greater for OA-LA than YA at 90 minutes (−512±477 pg·mL^-1^ vs. 174±182 pg·mL^-1^, *p*=0.045) and 180 minutes (−502±147 pg·mL^-1^ vs. −208±202 pg·mL^-1^, *p*=0.049), and lower than OA-HA at 60 minutes (−447±447 pg·mL^-1^ vs. −125±169 pg·mL^-1^, *p*=0.039). GLP-1 concentration was higher for OA-LA compared with YA at 180 minutes (5.00±4.71 pM vs. 1.07±2.83 pM, *p*=0.040). Net AUC for PYY response to feeding was greater for OA-LA compared with OA-HA (*p*=0.052). No differences were seen in subjective appetite. These observations in older adults exhibiting an anorexia of ageing phenotype suggest augmented anorexigenic responses of gut hormones to feeding may be causal mechanisms of anorexia of ageing.

## INTRODUCTION

Anorexia of ageing describes the age-related decline in appetite and food intake experienced in later life (Morley, 1997). A loss of appetite affects over 30% of community dwelling older adults (van den Broeke et al., 2018) and up to 60% of older adult hospital patients (Cox et al., 2020; Ray et al., 2014). Anorexia of ageing has been strongly implicated in malnutrition (Dent et al., 2019), which is associated with sarcopenia, frailty (Ligthart-Melis et al., 2020), and mortality (Söderström et al., 2017). The subsequent increased healthcare utilisation is substantial. Annual health and social care costs are estimated to be 3 times greater for undernourished older adults, compared with those adequately nourished (Russell & Elia, 2014). With an ageing global population, malnutrition in later life is an imposing challenge for current and future healthcare provisions.

The causes of anorexia of ageing are yet to be conclusively determined. It is likely a multifaceted phenomenon, including age-related changes in physiological and hedonic control, and societal factors (Cox et al., 2020). One proposed mechanism is a change in appetite-associated gut hormone secretion with age. A meta-analysis by Johnson et al. (2020) showed elevated concentrations of the anorexigenic hormones leptin, CKK, and PYY in older adults compared with younger adults. However, the effect of ageing on concentrations of other appetite-associated hormones, such as ghrelin and GLP-1 were less clear.

A potential reason for the remaining contention regarding age-related changes in gut hormones – and regarding other mechanisms of anorexia of ageing – relates to the common design of studies in this field. Typically, studies compare mechanisms of interest, such as gut hormone responses to feeding, between younger adults and older adults, with little consideration of the heterogeneity of the older adult cohort. Heterogeneity in eating behaviour (ter Borg et al., 2015) and nutritional needs of older adults (Krondl et al., 2008) is well-established, and has been identified as a challenge when attempting to identify relationships between participant characteristics and eating patterns or weight status (Hsiao et al., 2011). In addition, variance in gut hormone responses to feeding in older adults is often considerable (Johnson et al., 2020). Indeed, with the prevalence of anorexia of ageing in community dwelling older adults being around 30%, it is likely that study cohorts of older adults consist of some with impaired appetite and some with unimpaired, healthy appetite. Pooling both in the same study cohort likely masks some responses that are not a function of ageing but are exclusive to those with suppressed appetite. Consequently, hormonal dysregulation that may be causal of anorexia of ageing could be overlooked.

Identifying those with low appetite is challenging. The limitations of free-living, self-reported measures of habitual dietary intake are well-known (Ravelli & Schoeller, 2020; Saravia et al., 2022), especially in older adults where recall bias may be increased (Freedman et al., 2014; Park et al., 2018; Rhodes et al., 2019) and adherence to food diaries has been shown to be low (Rowland et al., 2018). Changes in body mass, indicating inadequate energy intake, are not always detected as only around 50% of people self-weigh regularly (Gavin et al., 2015; VanWormer et al., 2012) and access to weighing scales is limited for some cohorts of the population (Bramante et al., 2020). Questionnaires have been developed for assessing appetite, such as the Simplified Nutritional Appetite Questionnaire (SNAQ). This tool has proved a quick and simple way to identify individuals at risk of undernutrition, with validity having been shown in community-dwelling (Lau et al., 2020) and hospitalised patients (Kruizenga et al., 2005). However, there is contention over cut-off points for identifying low-appetite (Wilson et al., 2005; Lau et al., 2020) and conformation of criterion validity against an objective measure of eating behaviour or appetite is lacking.

Recently, we used a multi-criteria approach, including an objective, laboratory-measured assessment of energy intake at an *ad libitum* test meal, to identify older adults with low appetite. This model enabled us to observe differences in ghrelin metabolism between healthy-appetite older adults and low-appetite older adults (Holliday et al., in submission). Phenotyping older adults exhibiting anorexia of ageing in this way should facilitate the exploration of the mechanisms underpinning why some older adults experience low appetite and some do not.

The aim of this study was to determine gut hormone response to feeding in older adults with apparent healthy appetite and older adults exhibiting low appetite. We aimed to confirm our recent findings of differences in ghrelin response in a slightly extended sample, in combination with determining responses of anorexigenic hormones PYY and GLP-1. Comparing gut hormone responses to feeding between younger adults, older adults with a healthy appetite, and older adults with low appetite will shed light on changes in gut hormones that reflect normal ageing and those which may underpin the age-related decline in appetite and energy intake characteristic of anorexia of ageing. A secondary aim was to assess the appropriateness of our four-criteria method of phenotyping older adults with low appetite.

## METHOD

### Study Design

In a cross-sectional, observational study, responses of ghrelin, PYY, and GLP-1 to feeding were compared between younger adults (YA, aged 18 – 29 years), older adults (aged ≥ 65 years) with a healthy appetite (OA-HA), and older adults with low appetite (OA-LA). The study adhered to the ethical guidelines as outlined in the Declaration of Helsinki, and gained ethical approval from the Newcastle University Faculty of Medical Sciences Research Ethics Committee (LREC #: 2146/13433/2020).

### Participants

Fifteen non-obese, low-to-moderately active YA; and thirty non-obese, low-to-moderately active OA were recruited. Inclusion criteria were a score of < 3000 MET mins · week^-1^ on the International Physical Activity Questionnaire (IPAQ; Craig et al., 2003), body mass index (BMI) of < 30 kg·m^-2^ for YA and < 33 kg·m^-2^ for OA (such a BMI value is associated with increased risk of mortality in older adults (Winter et al., 2014)), non-smoker, not attempting to intentionally change bodyweight or composition, not taking medication likely to impact on appetite, and free from metabolic disease. OA were categorised as either exhibiting a healthy appetite (OA-HA) or exhibiting signs of low appetite (OA-LA). Low appetite was identified if two of four *a priori* criteria were met (Holliday et al., in submission). These were:

1. Low BMI (< 23 kg·m^-2^ (such a BMI value is associated with increased risk of mortality in older adults (Winter et al., 2014)).
2. Low habitual energy intake (<75% estimated total energy requirement (TER), as identified by the World Health Organisation as indicative of undernutrition) as measured by 24-hour dietary recall.
3. Low score (< 15) on the Simplified Nutritional Appetite Questionnaire (SNAQ; Lau et al., 2020).
4. A laboratory-measured *ad libitum* lunch intake of < 25% of estimated TER (based on a typical lunch energy intake of ∼27% of total energy intake in UK mid-life adults (Pot et al., 2014)).

Younger adults who met two of these four criteria (low BMI cut off of < 18.5 kg·m^-2^) were excluded.

### Enrolment and Familiarisation

Participants visited the Human Nutrition Suite at Newcastle University for enrolment and familiarisation. Informed written consent was obtained after the study procedures had been explained verbally and after any questions had been addressed. Height and weight were recorded, and habitual physical activity (IPAQ) and appetite (SNAQ) were assessed. An assessment of habitual daily food intake was obtained using the computerised, multiple-pass, 24-hour dietary recall system, Intake24 (Foster et al., 2019). Total daily energy requirement was estimated using the Mifflin-St Joer equation (Mifflin et al., 1990).

Participants were then familiarised with the test meals to be consumed on the trial visit. The breakfast meal was provided in full to ensure all participants could finish the entire portion in the standardised time of between five and six minutes. Those unable to consume the entire portion were excluded from the study. Palatability of the lunch meal was also confirmed by providing a small sample to taste.

### Experimental Procedures

Participants returned to the Human Nutrition Suite at Newcastle University within two weeks of the enrolment visit for the experimental trial. Participants were instructed to abstain from exercise, caffeine, and alcohol on the day before the experimental visit, and to consume a standardised, nutrient-balanced evening meal of beef hash, yoghurt and orange juice (691 kcal; 47% energy from carbohydrate, 29% fat, 23% protein) a minimum of 12 hours prior to arrival at the laboratory the following day. Participants arrived at the laboratory between 08:00 and 09:00, fasted but having drunk 300mL of water upon waking. On arrival, adherence to dietary and exercise controls was confirmed. Subjective appetite was then assessed using the visual analogue scale (VAS) method, before a cannula was inserted into the antecubital vein of the arm (time: t=0 minutes, see Figure 1). Ten minutes after cannulation (t=10 minutes), a fasted blood sample was obtained. At t=15 minutes, participants consumed a standardised breakfast test meal of porridge (made with whole milk) with natural yoghurt and honey. The energy content of the meal was 450kcal, with a macronutrient balance representative of UK dietary recommendations (50% carbohydrate, 15% protein, 35% fat). Preparation of the porridge breakfast meal, including cooling time, was identical for all participants, and all participants consumed the meal in between five and six minutes.

**Figure 1.**
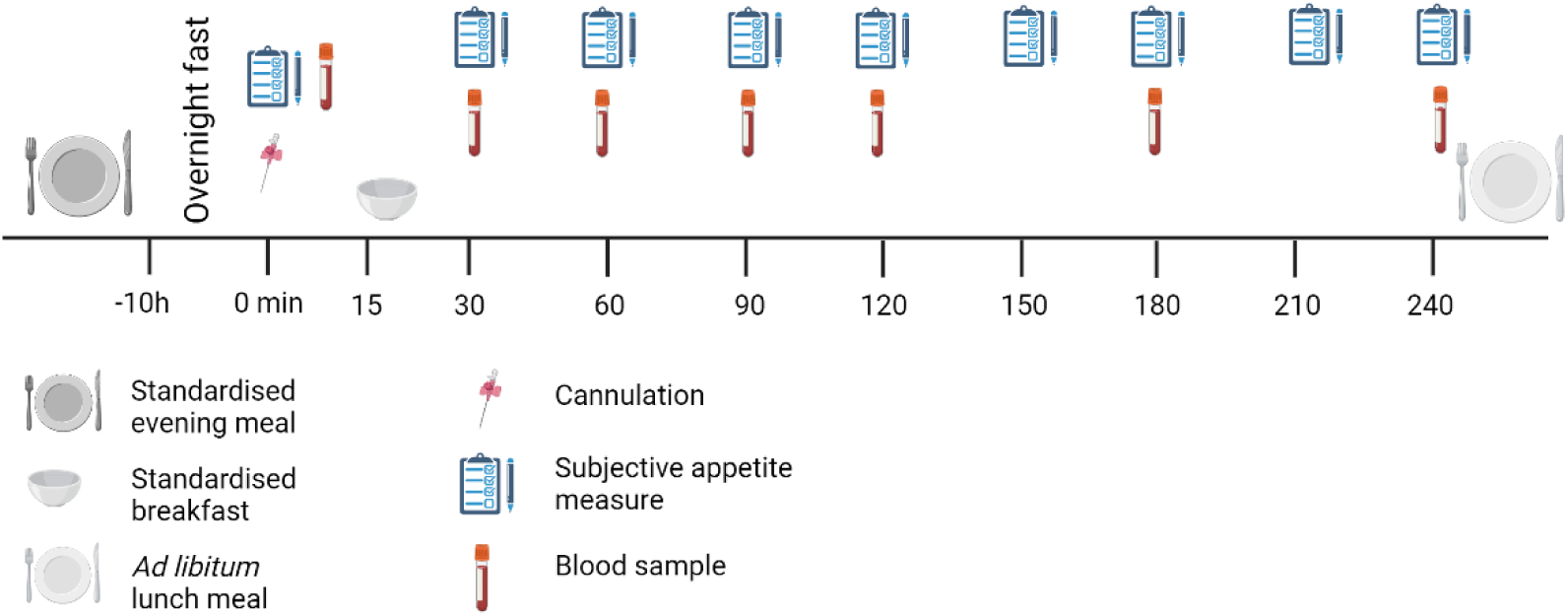
Study protocol.

At t=30 minutes, subjective appetite was measured and a second blood sample was obtained. Participants then rested for a further 210 minutes, with appetite measured every 30 minutes and blood samples obtained at t=60, 90, 120, 180 and 240 minutes (see Figure 1). During this period, participants remained seated and were free to read, watch television or use a laptop computer. Activity was monitored to ensure the avoidance of food cues in reading and viewing material.

At t=240 minutes, the cannula was removed and participants were provided with an *ad libitum* pasta-based lunch meal. Upon completion of the meal, the trial was complete.

### Outcome Measures

#### Plasma concentration of ghrelin, PYY and GLP-1

Blood was collected in EDTA-treated blood collection tubes for the measure of total PYY and GLP-1. For ghrelin, blood was collected in EDTA tubes pre-treated with AEBSF protease inhibitor at a concentration of 1g·mL^-1^ of whole blood (Deschaine & Leggio, 2020). Whole blood was centrifuged at 2000g and 4C for 15 minutes to separate plasma from cellular material. Plasma was aliquoted into 0.5mL sample cups and stored at −80ᵒC for later analysis. Plasma aliquots for the measure of ghrelin were treated with 0.2mL of 1M hydrochloric acid.

Total ghrelin was measured by enzyme-link immunosorbent assay (ELISA). Ghrelin was measured using commercially available kits (Human Ghrelin (total) ELISA kit, Merck Millipore). Sensitivity was 156 pg·mL^-1^. Coefficients of variation (CV) was 6.38%. Samples from 35 participants (11 YA, 11 OA-HA, 13 OA-LA) were measured for total GLP-1 and total PYY using in-house established radioimmunoassay (RIA) at Imperial College London (Kreymann et al., 1987, Adrian et al., 1985). Sensitivity and CV of RIA were 0.36 pg.mL^-1^, 4.43% and 2.885 pg.mL^-1^, 3.97% for GLP-1 and PYY respectively. Samples from 5 participants (1 YA, 2 OA-HA, 2 OA-LA) were measured by ELISA using commercially available kits (Human PYY (Total) ELISA kit and Multi Species GLP-1 Total ELISA, Merck Millipore) due to the unavailability of RIA labels. Sensitivity and CV were 1.5 pM, 6.95% and 1.4 pg.mL^-1^, 3.28% for GLP-1 and PYY respectively.

#### Subjective appetite

Measures of subjective appetite perceptions were obtained using the 4-item VAS method, assessing hunger, fulness, desire to eat and expected consumption (Flint et al., 2000). A composite score was calculated from the four items as: (hunger score + (100-fullnessscore) + desire to eat score + expected intake score) / 4 (Holliday & Blannin, 2017).

#### Ad libitum food intake

Food intake was assessed with a homogeneous pasta-based *ad libitum* test meal (Deighton et al., 2016). The meal was nutrient-balanced to align with UK dietary recommendations and consisted of pasta, Bolognese sauce and grated cheese, with added olive oil (energy density = 1.79 kcal·g^-1^. 50% energy from carbohydrate, 15% protein, 35% fat). Participants were instructed to eat until they felt “satisfyingly full.” To avoid a situation whereby an empty bowl provided a cue to stop eating prior to satiation, each bowl of pasta was replaced with a fresh, full bowl before the participant emptied the previous one. Food was consumed in isolation, with an avoidance of distractions and food cues, and with no time limit. Energy intake was calculated from the mass of food consumed and the known energy density of the meal.

### Statistical Analyses

All values are presented and mean ± SD (mean ± SEM in figures). Fasted ghrelin, PYY, and GLP-1 concentrations and lunch *ad libitum* EI (expressed as absolute intake and intake as a percentage of estimated TER) were compared between YA and all OA by independent samples t-tests. Differences between YA, OA-HA, and OA-LA were assessed by one-way analysis of variance (ANOVA). Ghrelin, PYY, and GLP-1 response to feeding was presented as change-from-baseline concentration. Differences between groups (between-subject factor) and over the trial period (within-subject factor) were assessed using a mixed-design analysis of variance (ANOVA). Net area-under-the-curve (nAUC) was calculated for each of these variables using the trapezium method. Differences in nAUC between YA and all OA was assessed by independent samples t-test, and between YA, OA-HA, and OA-LA were assessed by one-way ANOVA. Significant interactions and main effects were explored further using Bonferroni-corrected pairwise comparisons. Eta squared (η^2^) and partial η^2^ (η^2^) effect sizes were calculated for main effects and interactions, respectively, while Cohen’s *d* effect sizes were calculated for pairwise comparisons. Statistical significance was determined at an alpha level of 0.05.

To assess predictors of anorexigenic responses of gut hormones to feeding, z scores for AUC were calculated for ghrelin, PYY and GLP-1. The ghrelin Z score was inverted and a mean Z score for all three hormones was calculated. This was termed “anorexigenic response score”, with a higher value representing a more anorexigenic response. Backward elimination linear regression was conducted with anorexigenic response score as the outcome variable and the four variables included in the criteria to identify low appetite (BMI, SNAQ score, daily EI as percentage of TER, and *ad libitum* lunch EI) as predictors. At each step, the least significant variable (above a *p*-value threshold of 0.1) was eliminated from the model until remaining variables contributed independently to variance in the outcome measure. Principle component analysis for BMI, SNAQ score, daily EI as percentage of TER, and *ad libitum* lunch EI was attempted but aborted due to violations of sampling adequacy (Kaiser-Meyer-Olkin test = 0.414). All statistical analyses were conducted using Statistical Package for Social Sciences (SPSS).

An *a priori* power calculation was conducted to power the study to detect changes in line with previous studies which had observed differences in PYY and GLP-1 concentration between older and younger adults (Geizenaar et al., 2018; Geizenaar et al., 2020). With statistical power of 0.8 and an alpha value of 0.05, a sample of at least 12 participants per group was required to detect a large difference (*d* = 0.8).

## RESULTS

### Participant characteristics

The characteristics of all participants included in analyses, as grouped by age and appetite are shown in Table 1. Two younger adults were excluded as they met two of the four criteria for identifying low appetite and one younger adult withdrew due to lack of time. One older adult withdrew due to lack of time, and one was excluded due to difficulty with phlebotomy procedures. The older and young adult cohorts did not differ by BMI, weight, or physical activity, but SNAQ score was significantly lower for older adults (*p* = 0.009, *d* = 0.953).

**Table 1.**
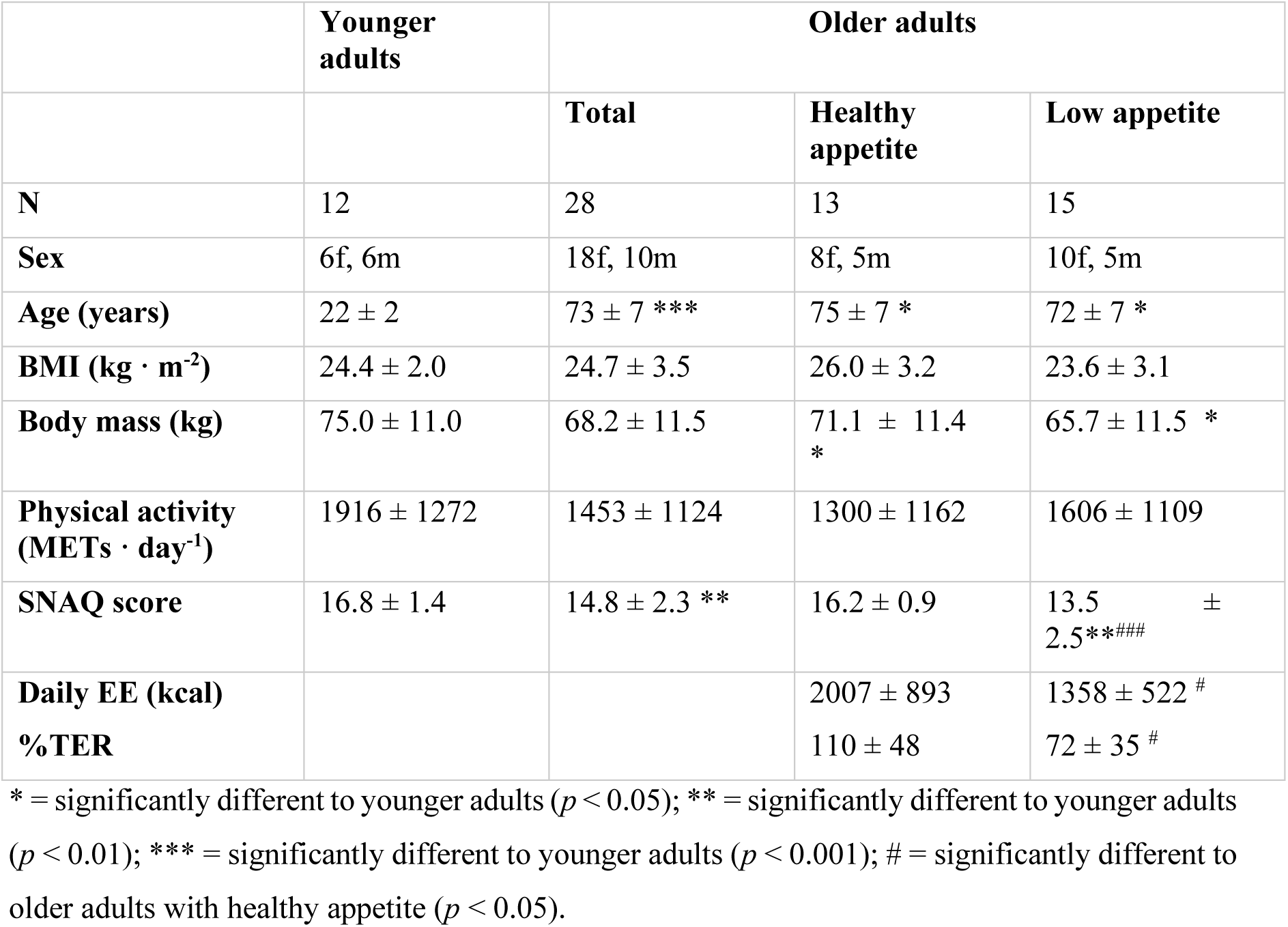
Participant characteristics for younger adults, all older adults, older adults with healthy appetite, and older adults with low appetite.

When comparing YA, OA-HA, and OA-LA, no significant differences were seen in body mass, BMI or physical activity (all *p* > 0.1). A group main effect for SNAQ score (*p* < 0.001, η^2^ = 0.405) was observed, with score being significantly lower in OA-LA compared with both YA (*p* < 0.001) and OA-HA (*p* = 0.001), but there was no difference between OA-HA and YA. Daily EI was lower in OA-LA compared with OA-HA (*p* = 0.046, *d* = 0.909), as was EI as percentage of TER (*p* = 0.041, *d* = 1.242). Age did not differ between OA-HA and OA-LA (*p* = 0.323).

### Energy intake

Absolute energy intake at the *ad libitum* lunch meal for YA, all OA, OA-HA and OA-LA is shown in Figures 2a. Energy intake was significantly greater for YA, compared with all OA (1108 ± 235 kcal vs. 532 ± 234 kcal, *p* < 0.001, *d* = 2.456). When comparing YA, OA-HA, and OA-LA, there was a significant group main effect (*p* < 0.001, η^2^ = 0.709). Post hoc pairwise comparisons revealed intake was significantly greater for YA (1108 ± 235 kcal), compared with both OA-HA (705 ± 207 kcal, *p* < 0.001, d = 1.820) and OA-LA (395 ± 150 kcal, *p* < 0.001, *d* = 3.617). Intake was also greater for OA-HA than OA-LA (*p* < 0.001, *d* = 1.713).

**Figure 2.**
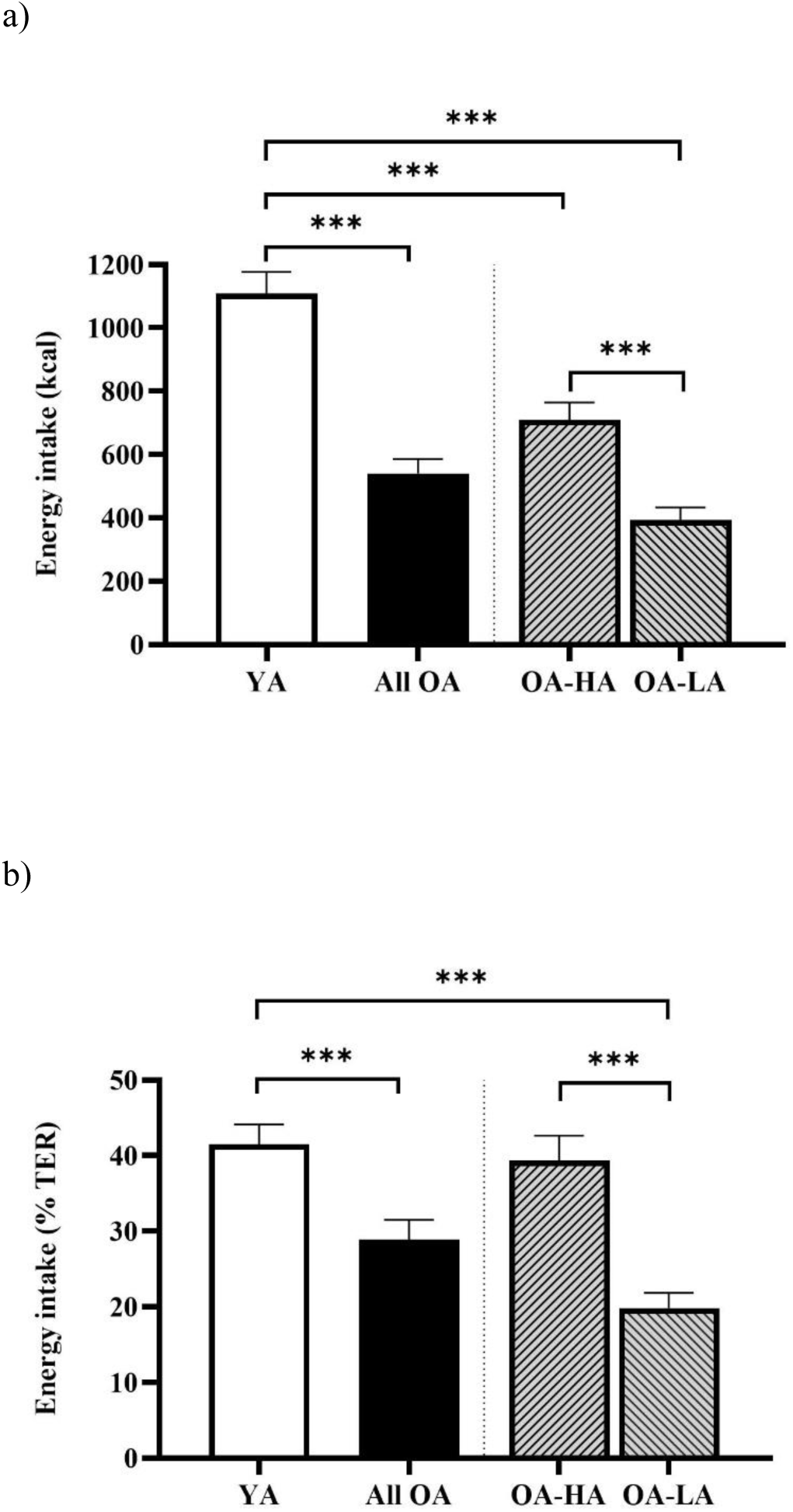
Mean ± SEM absolute lunch *ad libitum* energy intake (a) and lunch intake as a percentage of estimated TER (b) for YA, all OA, OA-HA, and OA-LA. *** = significant between-group difference, *p* ≤ 0.001.

When expressed relative to estimated TER (Figure 2b), energy intake as a percentage of TER was greater for YA compared with all OA (41.5 ± 9.2% vs. 27.6 ± 12.4%, *p* = 0.001, *d* = 1.207). When comparing YA, OA-HA, and OA-LA, there was a significant group main effect (*p* < 0.001, η^2^ = 0.561). Intake was lower for OA-LA (19.8 ± 7.7%) compared with both YA (41.5 ± 9.2%, *p* < 0.001, *d* = 2.558) and OA-HA (37.3 ± 10.0%, *p* < 0.001, *d* = 1.961). There was no difference between YA and OA-HA (*p* = 0.781).

### Fasted hormone concentrations

#### Ghrelin

Fasted plasma ghrelin concentration was significantly higher in OA compared with YA (1075 ± 654 pg·mL^-1^ vs. 636 ± 251 pg·mL^-1^, *p* = 0.007, *d* = 0.790, Figure 3a). When comparing YA, OA-HA, and OA-LA, there was a significant group main effect (*p* = 0.004, η^2^ = 0.284). Concentration was significantly higher in OA-LA (1332 ± 702 pg·mL^-1^) compared with both YA (636 ± 251 pg·mL^-1^, *p* = 0.005, *d* = 1.315) and OA-HA (771 ± 452 pg·mL^-1^, *p* = 0.034, *d* = 0.947).

**Figure 3.**
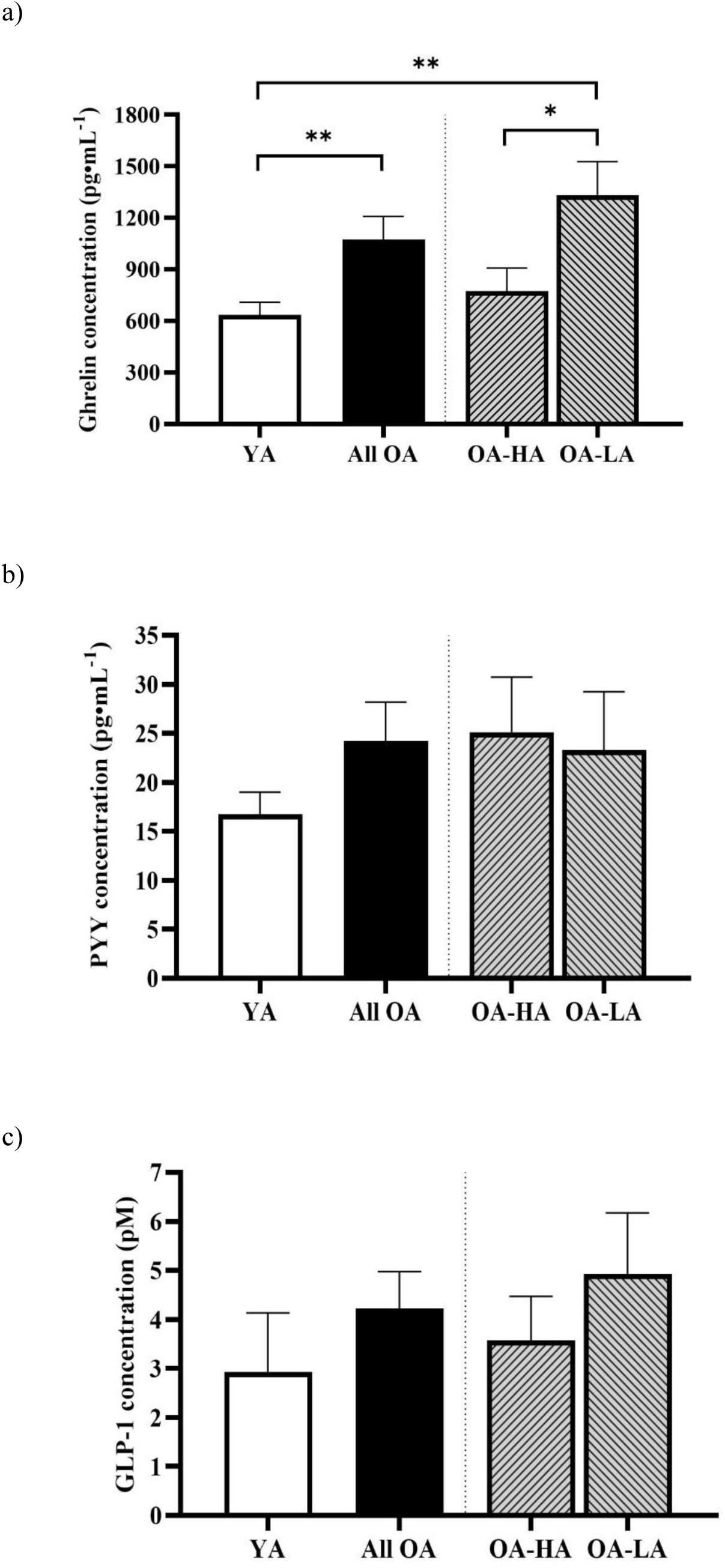
Mean ± SEM fasted concentrations of ghrelin (a), PYY (b), and GLP-1 (c). * = significantly different, *p* < 0.05. ** = significantly different, *p* < 0.01.

#### PYY

Fasted plasma PYY concentration did not differ between YA and all OA (16.75 ± 7.80 pg·mL^-1^ vs. 24.18 ± 21.63 pg·mL^-1^, *p* = 0.115, *d* = 0.395, Figure 3b), nor between YA, OA-HA, and OA-LA (16.75 ± 7.80 pg·mL^-1^ vs. 25.14 ± 20.87 pg·mL^-1^ vs. 23.29 ± 23.02 pg·mL^-1^, *p* = 0.512, η^2^ = 0.035).

#### GLP-1

Fasted plasma GLP-1 concentration did not differ between YA and all OA (2.93 ± 4.16 pM vs. 4.22 ± 3.93 pM, *p* = 0.357, *d* = 0.324, Figure 3c), nor between YA, OA-HA, and OA-LA (2.93 ± 4.16 pM vs. 3.57 ± 3.36 pM vs. 4.93 ± 4.50 pM, *p* = 0.450, η^2^ = 0.045).

### Hormone response to feeding

#### Ghrelin

The plasma ghrelin concentrations in response to the standardised breakfast meal are shown in Figure 4a and 4b. There was a significant group x time interaction when comparing YA and all OA (*p* = 0.049, η^2^ = 0.082, Figure 4a). Post hoc pairwise analysis showed a trend for a greater suppression in ghrelin concentration in OA at 180 min (−241.5 ± 320.0 pg·mL^-1^ vs. −48.0 ± 194.8 pg·mL^-1^, *p* = 0.064, *d* = 0.730). There was no group main effect (*p* = 0.268, η^2^ = 0.036) and nAUC did not differ between YA and all OA (−23899 ± 27733 pg·mL^-1^·240min^-1^ vs −48546 ± 62680 pg·mL^-1^·240min^-1^, *p* = 0.205, η^2^ = 0.047).

**Figure 4.**
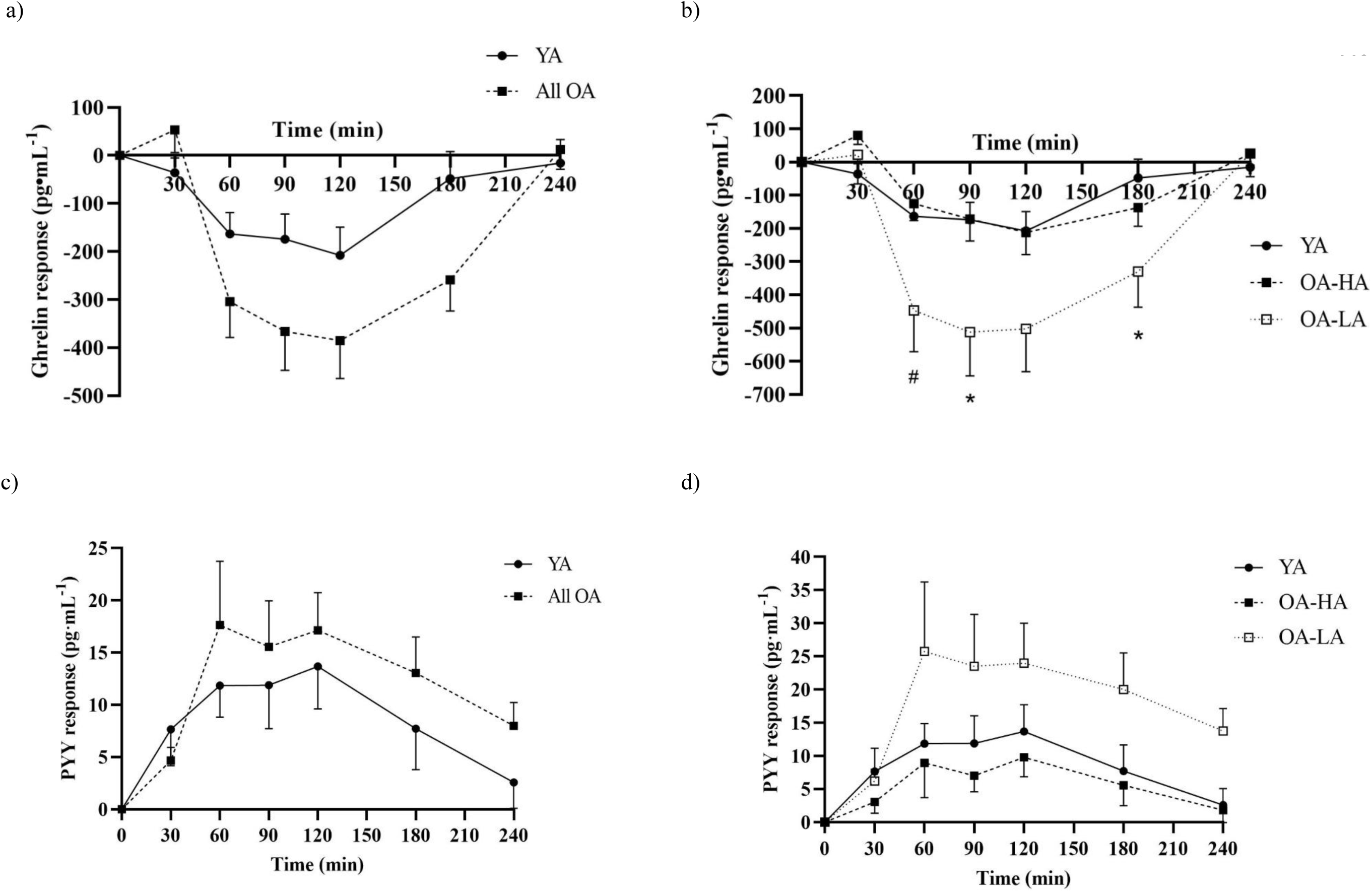
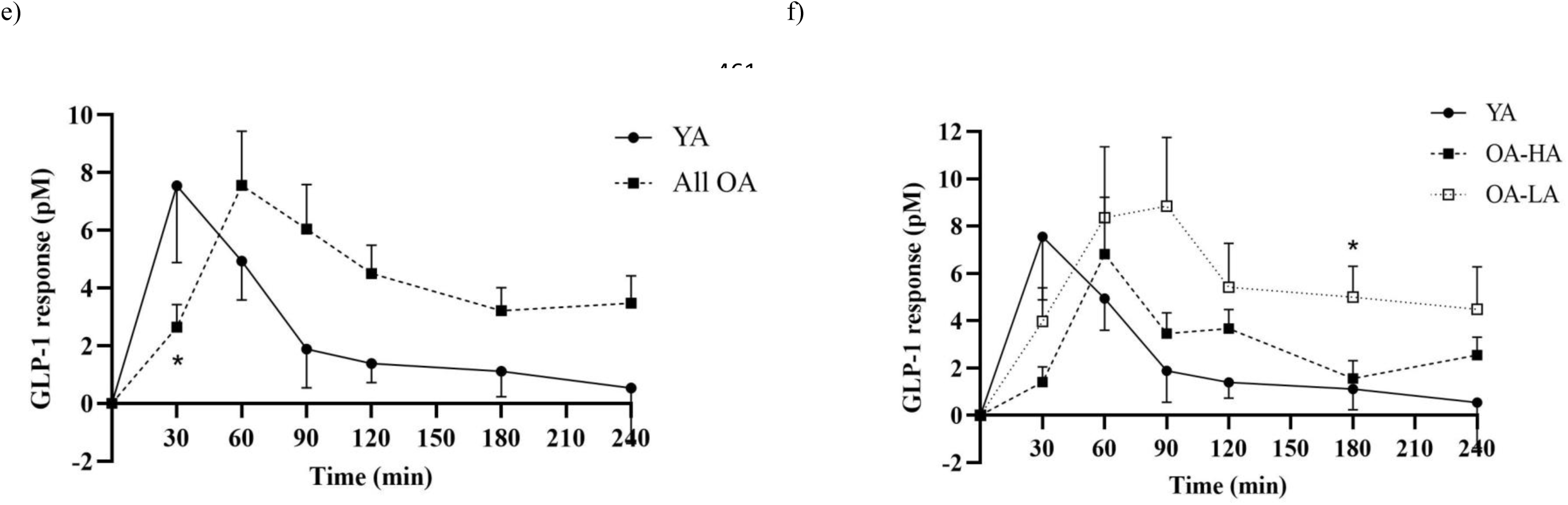
Mean ± SEM ghrelin (a and b), PYY (c and d), and GLP-1 (e and f) responses to feeding for YA (●, solid line) and all OA (▪, dashed line) (figures a, c, e) and for YA (●, solid line), OA-HA (▪, dashed line), and OA-LA (□, dotted line) (figures b, d, f). * = significantly different to YA, *p* < 0.05. # = significantly different to OA-HA, *p* < 0.05.

When comparing YA, OA-HA, and OA-LA, there was a significant group x time interaction (*p* = 0.010, η^2^ = 0.168, Figure 4b). Ghrelin concentration was lower in OA-LA compared with YA at 90 min (−512 ± 477 pg·mL^-1^ vs. 174 ± 182 pg·mL^-1^, *p* = 0.045, *d* = 1.900) and 180 (−502 ± 147 pg·mL^-1^ vs. −208 ± 202 pg·mL^-1^, *p* = 0.049, *d* =1.664), with a trend for a difference at 60 min and 120 min (both *p* < 0.01). Ghrelin concentration was significantly lower in OA-LA than OA-HA at 60 min (−447 ± 447 pg·mL^-1^ vs. −125 ± 169 pg·mL^-1^, *p* = 0.039, *d* = 0.953), with the difference at 90 min approaching significance (−512 ± 477 pg·mL^-1^ vs. 172 ± 219 pg·mL^-1^, *p* = 0.050, *d* = 1.843). The group main effect approached significance (*p* = 0.051, η^2^ = 0.165), with trend for a difference between OA-LA and OA-HA. There was a significant group main effect for nAUC (*p* = 0.046, η^2^= 0.170). Post hoc pairwise comparisons showed a trend for a more negative nAUC in OA-LA compared with YA (−69741 ± 74405 pg·mL^-1^·240min^-1^ vs −23899 ± 27733 pg·mL^-1^·240min^-1^, *p* = 0.095, *d* = 0.816).

#### PYY

The plasma PYY concentrations in response to the standardised breakfast meal are shown in Figure 4c and 4d. There was no significant group x time interaction when comparing YA with all OA (*p* = 0.564, η^2^ = 0.016, Figure 4c).There was also no group main effect (*p* = 0.507, η^2^ = 0.11) and no difference in nAUC between YA and OA (2097 ± 2314 pg·mL^-1^·240min^-1^ vs 2930 ± 3749 pg·mL^-1^·240min^-1^, *p* = 0.481, *d* = 0.244).

When comparing YA, OA-HA, and OA-LA, there was no significant group x time interaction (*p* = 0.450, η^2^ = 0.048, Figure 4d). The group main effect approached significance (*p* = 0.051, η^2^ = 0.145), with a trend for a difference between OA-LA and OA-HA (*p* = 0.058). There was a significant group main effect for nAUC (*p* = 0.045, η^2^ = 0.151), with the greater nAUC in OA-LA compared with OA-HA approaching significance (4357 ± 4662 pg·mL^-1^·240min^-1^ vs 1400 ± 1416 pg·mL^-1^·240min^-1^, *p* = 0.052, *d* = 0.858).

#### GLP-1

The plasma GLP-1 concentrations in response to the standardised breakfast meal are shown in Figure 4e and 4f. There was a significant group x time interaction when comparing YA and all OA (*p* = 0.003, η^2^ = 0.101, Figure 4e). There was a more immediate increase in GLP-1 at 30 mins in YA compared with OA (7.55 ± 9.24 pM vs. 2.64 ± 4.08 pM, *p* = 0.026, *d* = 0.687). However, GLP-1 remained elevated in OA, with a trend for a higher concentration at 120 min (4.51 ± 5.09 pM vs. 1.38 ± 2.30 pM, *p* = 0.05, *d* = 0.792). Net AUC was not significantly different between YA and OA (576 ± 663 pM·240min^-1^ vs. 987 ± 1012 pM·240min^-1^, *p* = 0.207, *d* = 0.446).

When comparing YA, OA-HA, and OA-LA, there was a significant group x time interaction (*p* = 0.041, η^2^_p_ = 0.096, Figure 4f). GLP-1 concentration remained higher in OA-LA compared with YA at 180 mins (5.00 ± 4.71 pM vs. 1.07 ± 2.83 pM, *p* = 0.040, *d* = 1.011), with a trend for a difference between at 120 min (5.41 ± 6.68 pM vs. 1.69 ± 2.65 pM, *p* = 0.099, *d* = 0.732). There was a trend for a difference in nAUC (*p* = 0.093, η^2^_p_ = 0.124).

#### Subjective appetite

When comparing YA with all OA, there was no difference in baseline subjective appetite score (67.4 ± 16.2mm vs. 60.2 ± 17.7mm, *p* = 0.235, *d* = 0.417. Figure 5a). When assessing appetite response to the standardised breakfast, as change-from-baseline, there was no significant group x time interaction (*p* = 0.155, η^2^ = 0.046), nor group main effect (*p* = 0.657; η^2^ = 0.005) for subjective appetite across the trial period. Net AUC did not differ between groups (*p* = 0.602, *d* = 0.181).

**Figure 5.**
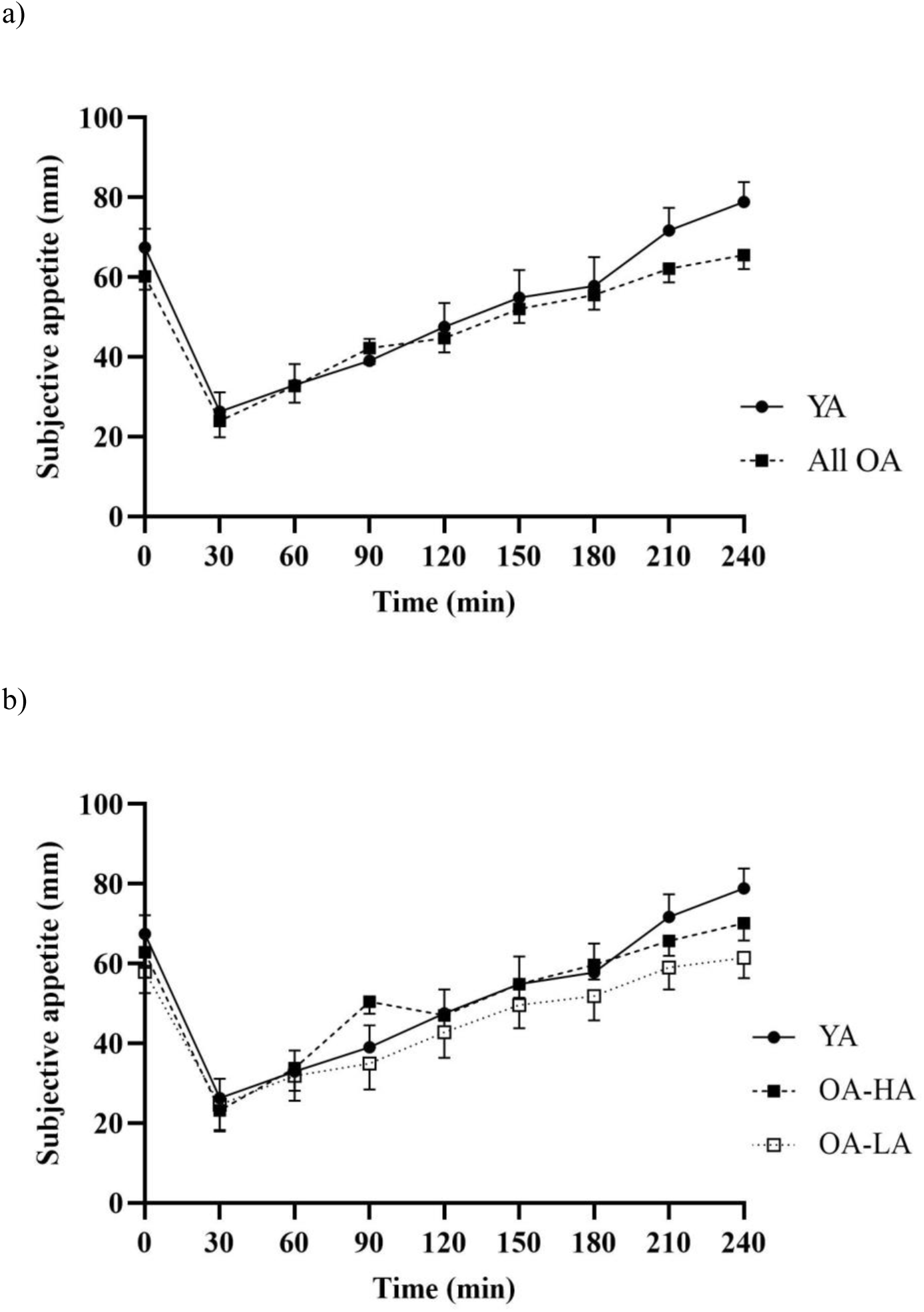
Mean ± SEM subjective appetite for YA vs. all OA (a), and YA vs. OA-HA vs. OA-LA (b).

When comparing YA, OA-HA, and OA-LA, there was no significant difference in baseline subjective appetite score (YA = 67.4 ± 16.2mm, OA-HA = 62.8 ± 14.0mm, OA-LA = 58.0 ± 20.6mm; *p* = 0.380, η^2^_p_ = 0.051. Figure 5b). There was also no significant group x time interaction (*p* = 0.169, η^2^_p_ = 0.080) nor group main effect (*p* = 0.896, η^2^ = 0.006) for subjective appetite response to the standardised breakfast. Net AUC did not differ between groups (*p* = 0.867, η^2^ = 0.008).

#### Regression analysis

The correlation matrix for the four predictor variables (BMI, daily energy intake as a percentage of TER, SNAQ score, and laboratory *ad libitum* lunch intake) and the outcome variable (anorexigenic response score) is shown in Table 2. SNAQ score and *ad libitum* lunch EI were significantly, negatively associated with anorexigenic response score. SNAQ score and *ad libitum* lunch intake were significantly correlated with one another.

**Table 2.** Correlation matrix for predictor variables BMI, daily EI as a percentage of TER, SNAQ score, and *ad libitum* lunch and outcome variable anorexigenic response score.

|  | Anorexigenic response score | BMI | Daily EI | SNAQ score | *Ad libitum* EI |
| --- | --- | --- | --- | --- | --- |
| Anorexigenic response score |  | -0.158 | -0.350 | -0.647 *** | -0.419 * |
| BMI |  |  | -0.358 | 0.198 | 0.162 |
| Daily EI |  |  |  | 0.168 | 0.235 |
| SNAQ score |  |  |  |  | 0.470 ** |
| *Ad libitum* EI |  |  |  |  |  |

\*\*\* = $p < 0.001$ , \*\* = $p < 0.01$ , \* = $p < 0.05$ .

**Table 3.** Backward elimination regression analyses of SNAQ score, daily EI, *ad libitum* lunch EI, and BMI as predictors of anorexigenic response of gut hormones.

| Model | R<sup>2</sup> | Adj R<sup>2</sup> | Predictors | β | p |
| --- | --- | --- | --- | --- | --- |
| 1 | 0.509 | 0.386 | SNAQ score | -0.538 | **0.012** |
|  |  |  | Daily EI | -0.247 | 0.258 |
|  |  |  | *Ad libitum* EI | -0.154 | 0.478 |
|  |  |  | BMI | -0.065 | 0.771 |
| 2 | 0.506 | 0.420 | SNAQ score | -0.549 | **0.008** |
|  |  |  | Daily EI | -0.215 | 0.239 |
|  |  |  | *Ad libitum* EI | -0.181 | 0.341 |
| 3 | 0.478 | 0.419 | SNAQ score | -0.605 | **0.003** |
|  |  |  | Daily EI | -0.248 | **0.168** |
| 4 | 0.419 | 0.388 | SNAQ score | -0.647 | **0.002** |

Regression analysis showed that a model containing all four predictor variables was a significant predictor of gut hormone anorexigenic response, with variance in these variables explaining 51% of variance in gut hormones response. In the model containing all four predictors, the only significant predictor or gut hormone anorexigenic response was SNAQ score (β = −0.538, *p* = 0.012).

Three backward eliminations were performed, producing a total of four models. The decrease in R^2^ with each elimination was not significant. The model with the greatest predictive power, as denoted by the adjusted R^2^ value, was the model containing SNAQ score, daily EI and *ad libitum* lunch EI (adjusted R^2^ = 0.420).

## DISCUSSION

Our primary aim was to determine the ghrelin, PYY and GLP-1 responses to feeding in both older adults with unimpaired, healthy appetite and older adults with low appetite. Our novel findings show augmented anorexigenic gut hormone responses to feeding in older adults identified as having low appetite, but not in older adults with a healthy appetite. Suppression of the hunger hormone ghrelin was greater in older adults with low appetite, compared with both younger adults and older adults with healthy appetite. Increases in postprandial plasma concentration of the satiety hormones PYY and GLP-1 were greater and more enduring in older adults with low appetite, compared with younger adults. This was not observed in older adults with healthy appetite. Therefore, we propose that augmented anorexigenic responses of gut hormones to feeding is not a function of ageing *per se*, but instead may be a causal mechanism of anorexia of ageing.

Our approach of identifying older adults with low appetite allowed for comparisons between all older adults and young adults, and between older adults with low appetite, older adults with healthy appetite, and younger adults. When making comparisons purely on age, we observed greater postprandial increases in GLP-1 in older adults than younger adults, and non-significant greater postprandial responses of ghrelin and PYY. When comparing older adults with low appetite, older adults with healthy appetite, and young adults, it was revealed that these apparent age-related differences in gut hormone concentrations were driven by responses exclusively seen in older adults with low appetite.

Previous studies had evidenced age-related differences in postprandial concentration of ghrelin (di Francesco et al., 2008; Nass et al., 2014), PYY (Giezenaar et al., 2018a), and GLP-1 (Giezenaar et al., 2018b; Giezenaar et al., 2020). Our data indicate that such differences are not functions of ageing *per se* but are unique and specific to those with impaired appetite. Other studies have shown no difference in postprandial ghrelin (Bauer et al., 2010; Bertoli et al., 2006; Giezenaar et al., 2018a; Giezenaar et al., 2018b), PYY (di Francesco et al., 2005; MacIntosh et al., 1999) and GLP-1 (MacIntosh et al., 1999; MacIntosh et al., 2001; Trahair et al., 2012; Herpich et al., 2022) concentrations between older and younger adults. It is possible that these studies failed to observed differences due to the older adult cohort largely consisting of non-appetite suppressed older adults. Recruiting older adult study cohorts heterogeneous in appetite regulation, perceptions and eating behaviour could mask dysregulation of gut hormone responses exclusive to those with low appetite.

An amplified response of gut hormones to feeding is indicative of hypersensitivity of the gut to nutrient delivery. Gut hormones are secreted from specialised enteroendocrine cells of the GI tract in response to the sensing of nutrients or to changes in nutrient status. PYY and GLP-1 are secreted from enteroendocrine L-cells of the small and large intestine (Eissele et al., 1992; Sjölund et al., 1983), while ghrelin is secreted from X/A cells in the epithelium of the stomach (Date et al., 2000). Secretion is regulated by the sensing of nutrients by nutrient receptors and transporters, and the subsequent activation of intracellular signalling pathways. A hypersecretory response to feeding, as observed in older adults with low appetite, would suggest upregulation, or dysregulation, of nutrient sensing or cellular signalling. As such, we further propose that augmented anorexigenic gut hormones response to feeding may be a result of hypersensitivity of the gut to nutrients, and this hypersensitivity may be a causal mechanism of anorexia of ageing.

The secondary aim was to assess the appropriateness of our approach to phenotyping older adults with low appetite. This was required for the comparison of gut hormone responses between older adults with a healthy appetite and older adults with low appetite in the present study, and an effective approach to phenotyping those with anorexia of ageing could prove beneficial for future research in this field. We adopted a four-criteria classification based on BMI, habitual daily energy intake, SNAQ score, and an objective, laboratory-measured *ad libitum* lunch energy intake. This approach has recently been deployed to determine differences in ghrelin metabolism (Holliday et al., in review). The regression analyses of the present study support the appropriateness of this classification model for identifying those with low appetite and phenotyping anorexia of ageing. Variance in the four criteria explained 51% of variance in gut hormone response in older adults. SNAQ score was the strongest individual predictor of gut hormone response, which supports the application of the SNAQ for identifying community-dwelling older adults with low appetite (Lau et al., 2020). The model with the strongest predictive power, however, included SNAQ score, habitual daily energy intake, and *ad libitum* lunch energy intake. This evidences the beneficial inclusion of an objective energy intake measure for identifying appetite. The inclusion of BMI to the model provided little additional predictive power. This is perhaps not surprising, as BMI appears not to be associated with protein-energy malnutrition in older adults (van der Pols-Vijlbrief et al., 2014), and between 20 and 35% of older adults with a BMI of greater than 25 kg·m^-2^ are at risk of undernutrition (Klee Oehlschlaeger et al., 2014; Özkaya & Gürbüz, 2019; Sulmont-Rossé et al., 2022).

Although the present study determined postprandial responses of ghrelin, PYY, and GLP-1, other gut hormones may be of interest. We did not measure cholecystokinin (CCK) or gastric inhibitory polypeptide (GIP). As there is evidence to suggest both hormones exhibit greater responses to feeding in older adults than younger adults (Giezenaar et al., 2018a; Giezenaar et al., 2018b; Johnson et al., 2020), it would have been interesting to confirm if such responses were also specific to those with low appetite. The effects of feeding on other gut hormones, such as pancreatic polypeptide (PP) and oxyntomodulin, have yet to be determined in older adults (Johnson et al., 2020). Further research is required to allow a more complete understanding of age-related changes in gut response to nutrients, and how such changes impact upon the appetite and eating behaviour of older adults.

## CONCLUSION

This is the first study to demonstrate that augmented anorexigenic responses of gut hormones to feeding are observed in older adults with low appetite but not in older adults with a healthy appetite. This highlights two different phenotypes of appetite regulation and response in older adults. As such, we propose that amplified gut hormone response, resulting from gut hypersensitivity to nutrients, may be a causal mechanism of anorexia of ageing. Future research is warranted to explore the presence of nutrient sensing and signalling dysregulation in appetite-suppressed older adults.

## ACKNOWLEDGEMENTS

We would like to thank our participants for taking part in the study.

## AUTHOR CONTRIBUTIONS

A.H., A.D., B.C., & G.F. conceived the research question. A.H., D.R.C., designed the study. A.H. and J.W. collected data. A.H. and A.D. conducted data analyses. A.H. wrote the manuscript. A.D V.C., B.C., D.R.C., & G.F. edited the manuscript. All authors approved the final version.

## FUNDING

This work was supported by a Wellcome Trust Institutional Strategic Support Fund award (recipient: A.H.) and an Ageing and Nutrient Sensing Network Early Career Researcher award (recipient: A.D.)

## CONFLICT OF INTEREST

The authors declare no conflicts of interest.

## ABBREVIATIONS

AEBSF: 4-(2-aminoethyl)benzenesulfonyl fluoride hydrochloride
ANOVA: Analysis of variance
nAUC: Net area under the curve
CCK: Cholecystokinin
EDTA: Ethylenediaminetetraacetic acid
ELISA: Enzyme-linked immunosorbent assay
HA-OA: Healthy appetite older adults
IPAQ: International Physical Activity Questionnaire
LA-OA: Low appetite older adults
METs: Metabolic equivalents
OA: Older adults
PP: Pancreatic polypeptide
PYY: Peptide tyrosine-tyrosine
SNAQ: Simplified Nutritional Appetite Questionnaire
TER: Total energy requirements
VAS: Visual analogue scale
YA: Young adults

## Notes

### Competing Interest Statement

The authors have declared no competing interest.

